# iGEM comes of age: trends in its research output

**DOI:** 10.1101/2021.01.21.424716

**Authors:** Ashwin K. Jainarayanan, Anastasios Galanis, Amatullah Mustafa Nakara, Guilherme E. Kundlatsch, Roger Rubio-Sánchez

## Abstract

The international Genetically Engineered Machine (iGEM) is an educational benchmark in synthetic biology. Eighteen years after its inception, it has also catalysed the infusion of synthetic biology with interdisciplinary fundamental and translational research as well as with inspired young scientists. Here, we communicate a quantitative analysis of compiled published work associated to iGEM projects, highlighting trends in their dissemination and versatility. As iGEM comes of age, we anticipate it will continue to revolutionise, alongside SynBio, several disciplines of science and industries through the development of synthetic biological systems towards a sustainable future.

---

Synthetic biology (SynBio) has emerged over the past two decades as the systematic and rational engineering of biological systems,^1^ aiming to impart genetically-engineered biological devices with novel functionalities^2^ or aspiring to instill life-like behaviours in artificial entities from the bottom-up.^3^ Nowadays, SynBio is an ever-growing field with potential to deliver world-class research and applications, alongside a budding multi-billion dollar industrial landscape.^4^

The international Genetically Engineered Machine (iGEM) competition is a platform in which high school, undergraduate, and overgraduate students follow engineering-based “design and test” approaches to tackle global issues through solutions centred around synthetic biological systems.^5^ Its highly interdisciplinary nature results in annually bringing together people from various fields of science, spanning from engineering to biological, physico-chemical and computer to social sciences. This, as shown schematically in Fig. 1a, not only fosters the development of WetLab and DryLab skills, but also provides students with a unique multifaceted experience to engage with, for example, notions around science communication and outreach, ethics and biosafety,^6^ policy making, and scientific responsibility.^7^

**Figure 1.**
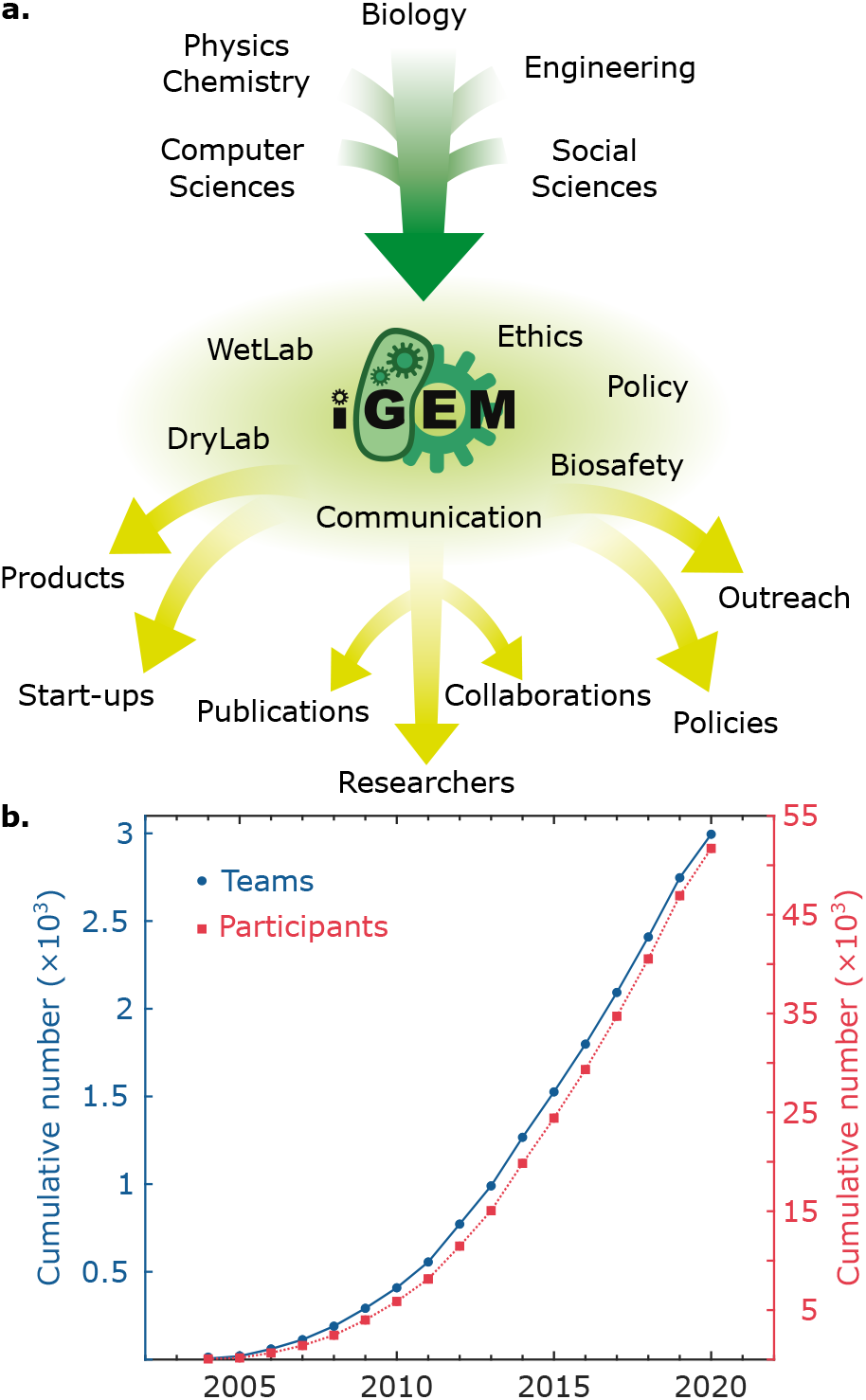
iGEM fuels Synthetic Biology in an evergrowing fashion. **a.** Schematic highlighting the interdisciplinarity of iGEM, a platform where scientists from several disciplines converge to develop synthetic biological systems aimed at tackling global issues. Participants engage with Wet and DryLab activities as well as with science communication and outreach, ethics and biosafety, and policy making, to name a few. In view of this multifaceted experience, its output includes – beyond the educational role – academic publications and young researchers. **b.** The community assembled by the annual competition is ever-increasing, as conveyed with a plot of the cumulative participation (teams in blue circles and participants in red squares) per year.

Indeed, aside from its educational role in the field,^8^ iGEM is also well-known as a venue for community building.^9^ Since its inception, > 50,000 alumni have participated associated to ~ 3000 teams, as collated in Fig. 1b. The event welcomes a nearly-increasing amount of teams each year (see Fig. S1), with ~ 250 of them during its latest edition. Impor-tantly, iGEM and SynBio have not only grown and matured concomintantly,^10^ but also in a cooperative fashion. In fact, the competition has fueled SynBio not only through innovative and visionary ideas and projects, some of which have found pathways to becoming start-ups and companies within the industrial sector, but also in the form of inspired young scientists eager to further the field *via* research and education.11 As iGEM alumni^i^, we reflect on our iGEM experiences and acknowledge the outstanding creativity, relevance, and impact underlining the projects within the competition.

In this communication, we investigate the academic output of iGEM projects whose scientific rigour, novelty, and applicability have been further disseminated in the form of publications. Our methodology, described schematically in the Supporting Information (see Fig. S2), harnessed a custom-built web scraping Python pipeline with which we identified and mapped preprint and peer-reviewed publications associated to iGEM projects; the latter was achieved by correlating available team and project information in www.igem.org with mined data containing the word *iGEM.* Matching publications were further analysed to extract quantitative metrics. Note that our strategy does not yet include publications associated to projects that recently participated in the 2020 edition.

In that sense, we identified that 109 manuscripts stemming from iGEM projects have been published since the competition was inaugurated in the early 2000s, as shown in Fig. 2a graphically. Interestingly, we observed sharp increases in publication numbers in 2014, and then later in 2018. Some of such observations, highlighted as peaks in Fig. S3 containing the non-cumulative publication trends, correlate in time with special issues and partnerships with journals (e.g. ACS SynBio iGEM 2013 Letters and PLOS iGEM Collections 2016, 2017), which were put in place with the aim of increasing the access to publishing project results within the iGEM community.

**Figure 2.**
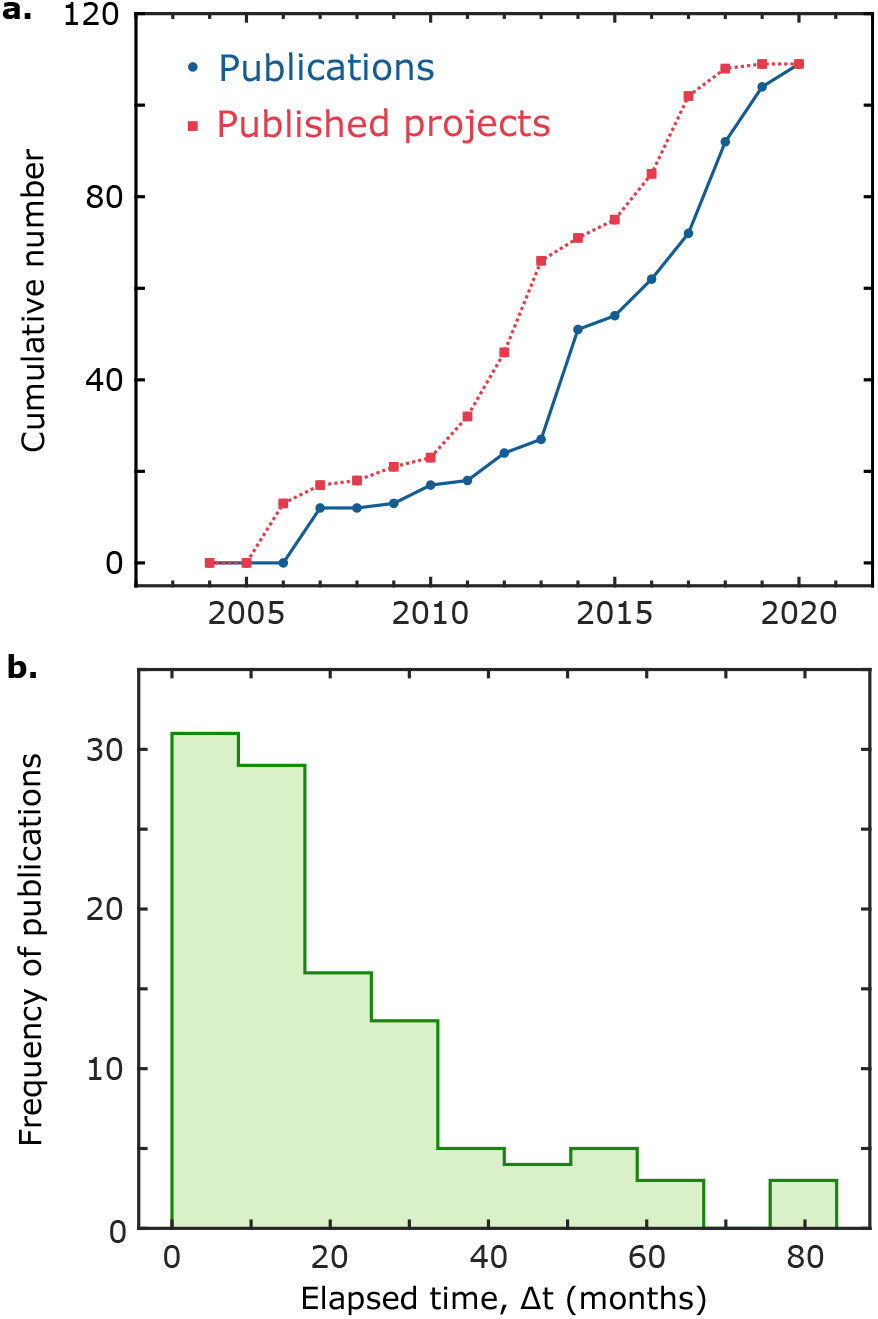
Trends in the dissemination of iGEM projects in the form of publications. Numerous iGEM projects have attained a publication stage, which warranted a “maturation time” after the iGEM season concluded. **a.** The frequency of publication of iGEM-associated projects over time, as conveyed by a plot of the cumulative number of publications (blue circles) and the cumulative number of published teams (red squares). One can readily observe a “phase difference” between the two curves. **b.** Elapsed time for publication, as conveyed by a histogram of ∆t for published teams, defined as the absolute difference between the publication date and that of the respective competition, set to the 31^st^ of October of the corresponding year.

Moreover, we note that when the publication numbers trend (blue) is compared to a curve outlining the number of published teams (red) in Fig. 2a, they simply appear to be “out-of-phase”. The latter hints at a substantial, yet consistent, time lag between the competition the projects participated in and their subsequent publication. To quantitatively gain insights from this “phase difference”, we defined an elapsed time for publication, ∆t, as the absolute difference (in months) between the date of publication and that of the competition the team was part of (set to 31^st^ of October of the corresponding year, since the season culminates with the Giant Jamboree annually taking place usually at the end of October). In turn, the computed *∆t* frequencies, presented in Fig. 2b as a histogram, readily showcase the distinct stages projects were at the time of the competition, where most of them required from 8 to 18 months to attain their publication. Interestingly, we also identified some cases where the project results were initially disseminated as a preprint, and published at a later date in a peer-reviewed journal (collated in Table S1 in the Supplementary Information).

Our main findings, summarised in Fig. 2, highlight the impact and novelty of iGEM projects, which presumably establish interesting and promising preliminary results by the end of the competition season (thoroughly documented in their Wiki entries). Nonetheless, projects appear to still warrant more time to “mature” and converge to a concise and well-rounded contribution to the field in the form of publications. In fact, we argue that in some instances, the judging process inherent to the competition might serve as an initial peer-review process, in which teams receive feedback that highlights the strengths of their projects as well as pointing out areas that may need more development.

Finally, the projects, their topics, and achievements span several disciplines, industries and applications. We thus classified published projects into categories, as determined from the track teams competed in. However, in some instances where teams did not have an assigned track (e.g. projects that competed before 2009), we simply categorised them in view of their topic/application to one of the available tracks. Figure 3 presents graphically the distribution of the areas published teams have contributed to over the years.

**Figure 3.**
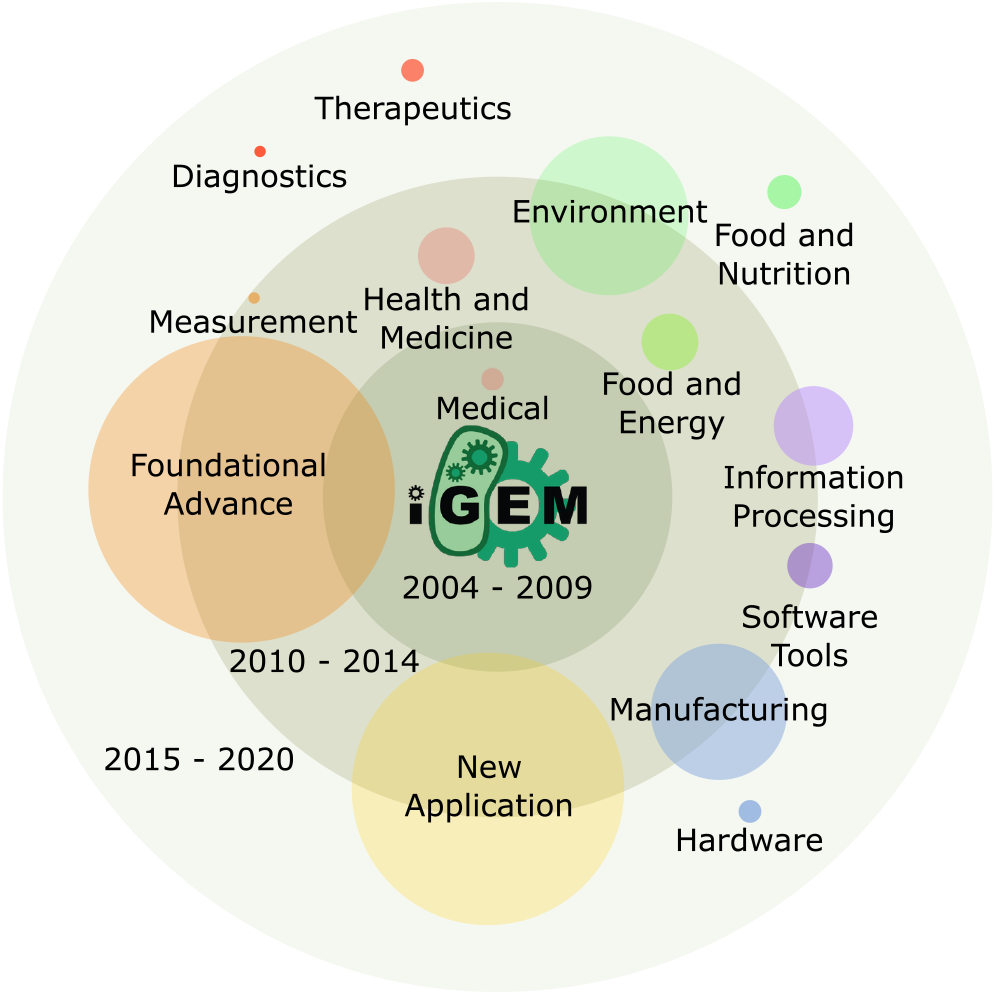
Published iGEM projects further a wide range of disciplines. iGEM-derived publications, centred around synthetic biological systems, address a breadth of fields and industries. Their versatility is showcased by a bubble chart, which highlights the distinct broad themes (categories) of published iGEM projects over time, as determined by the track each of them participated in. The radii of the bubbles are a proxy for the number of published teams within each category. Concentric circles represent time periods, as denoted by the labels in each of them, which aid in visualising the incidence of publications in categories as well as the evolution of the broad topics encompassed within the competition.

Here, the size of each bubble is a proxy for the number of teams/publications in each category. In addition, their location within the concentric rings roughly indicates their temporal publication incidence, while similar colour schemes exemplify the evolution of the broad topics encompassed within the iGEM competition.

Importantly, it can be readily observed that each of these projects has furthered the field in different disciplines. Moreover, the chart also aids in highlighting the versatility of synthetic biology as a means of addressing global issues. The proposals students have put forward for iGEM throughout the years have relevance or applicability ranging, for instance, from bio-medicine (health, diagnostics/therapeutics) to the environmental (energy, food and nutrition) and manufacturing (hardware) landscapes. Remarkably, iGEM projects in general, and not only those that have been published, as shown graphically in Supplementary Fig. S4, showcase the diversification over time that teams, alongside the competition and the field, have undergone to adapt and develop world-class solutions that tackle a breadth of social and scientific necessities.

In summary, we present a recapitulation of the academic output, in the form of publications, deriving from the iGEM competition. Having mined and quantified publications with roots in iGEM projects, we communicate trends in their dissemination (Fig. 2) and versatility (Fig. 3), which further highlight the role of the competition within SynBio. Notably, we acknowledge that our strategy may have not found other published, yet still iGEM-associated, work due to computational limitations or because these manuscripts do not have in their text *iGEM* as a word, which may result in an underestimation of the real publication numbers. We, thus, report – to the best of our knowledge – a compilation of the identified publications (see Supplementary File 1). Certainly, published iGEM work is a matter of broad interest to students, alumni and primary investigators. In fact, it has sprung initiatives such as partnerships with journals (as discussed above), but also student-led proposals resulting, for example, in the release of The Unofficial iGEM Proceedings Journal 2020, spearheaded by team iGEM Maastricht, in which original research pertaining to some 2020 projects was made available. This effort further exemplifies the relevance of accessible publishing venues and strategies aimed at globally delivering increased visibility of iGEM-associated research, as well as the broader need for barrier-free access to publishing, since the current regional publication landscape (see Fig. S5) shows trends seemingly favouring Europe alongside US and Canada.

Another important contribution – and example of the iGEM-derived academic output, yet not considered in our data set – comes from the interlaboratory study, which was aimed at standardising the calibration of cell density determination in liquid media.^12,13^ The latter promises to become a stepping stone for the engineering of reproducible synthetic biological entities.^14^ In this case, the authors use low-cost routes to determine cell numbers in a study with experimental contributions from ~ 260 investigators that spanned several iGEM teams, generations, and countries.

Furthermore, as iGEM ventures into its adulthood, in concert with SynBio commencing its third decade, it aims to deliver at least 10% of its projects as a readily-applicable real-world solution^15^ aligned to the Sustainable Development Goals (SDGs).^16^ We note that, as quantified above, ~ 4% of iGEM projects have been published either as a preprint or in a peer-reviewed venue. Although these observations are inspiring, they highlight the long way forward to harnessing the untapped potential that iGEM projects have. An interesting notion to spark, potentially as part of future competitions, would be to revive those projects that showed promise but were not further pursued, possibly due to lack of resources and/or team disbanding once the iGEM season concluded.

Finally, we argue that the percentage computed above is further increased by some other projects whose applicability enabled their transition to the commercial and/or start-up ecosystem, as identified in the Discovery & Insights 2020 Report prepared by the Entrepreneurship Program and Innovation Community (EPIC), with ~ 150 companies having connections to iGEM. We thus envision that the iGEM competition, its new era, and its ever-increasing community will continue to fuel the SynBio field and community to tackle the challenges ahead towards an equitable, inclusive, and sustainable future.

## Acknowledgement

The authors thank Athira Sreejith and Sourav Suresh (IISER Mohali) for support with literature mining and data curation, Gwyn Uttmark (Stanford) for feedback on the manuscript, and the rest of the Academia & Research Committee, along with the After iGEM team and iGEM Foundation, for useful conversations, insights, and support that helped in shaping this work and manuscript.

## Supporting Information Available

Supporting Information is available and includes:

- Supplementary Figures
- Supplementary File 1: Data sets, filtered with the analysis pipeline, culminating in the list of verified publications.
- Supplementary File 2: Processed data sets.

## Footnotes

i A.K.J. was part of the competition as student leader (2017, 2018) and advisor (2019); A.G. and A.M.N. participated as students (2019); G.E.K. was a student (2014) and advisor (2016); R.R.S. has been a student leader (2015), instructor (2016, 2017), andjudge (2019, 2020).

