## Supporting Information for "iGEM comes of age: trends in its research output"

### Supporting Information
### iGEM comes of age: trends in its research output.

Ashwin K. Jainarayanan,[*,†,‡] Anastasios Galanis,[†,¶] Amatullah Mustafa Nakara,[†] Guilherme E. Kundlatsch,[†,§] and Roger Rubio-Sánchez[*,†,‖]

†*Academia & Research Committee, After iGEM Team, iGEM Foundation, 45 Prospect Street, Cambridge, MA 02139, USA*

‡*The Kennedy Institute of Rheumatology, University of Oxford, Roosevelt Dr, Headington, Oxford OX3 7FY, United Kingdom*

¶*Institute for Fundamental Biomedical Research, Biomedical Sciences Research Center "Alexander Fleming", 16672 Vari, Greece*

§*School of Pharmaceutical Sciences, São Paulo State University (UNESP), Rodovia Araraquara-Jau Km 1, 14800-903 Araraquara, Brazil*

‖*Biological and Soft Systems, Cavendish Laboratory, University of Cambridge, JJ Thomson Avenue, Cambridge CB3 0HE, United Kingdom*

### Supplementary Figures

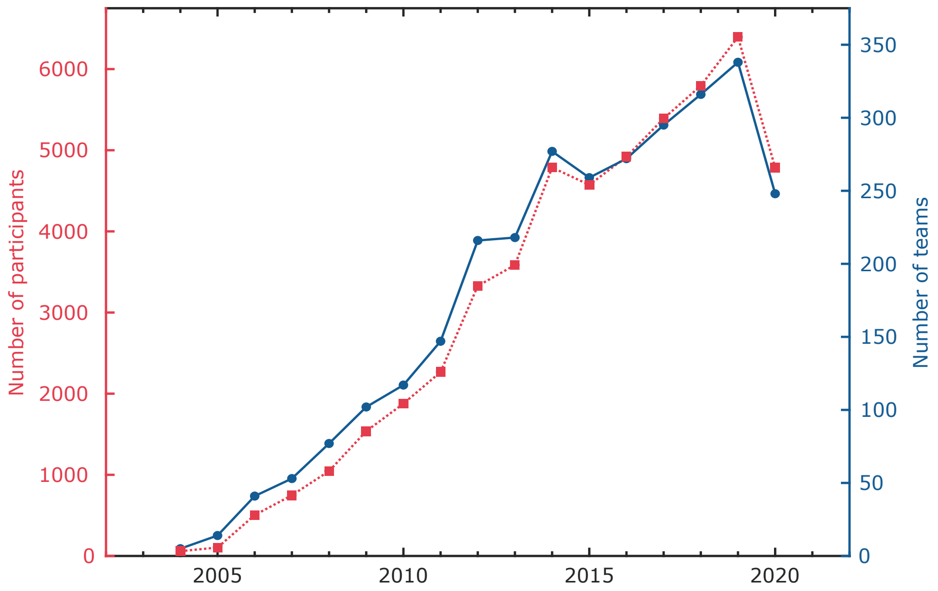


**Figure S1: iGEM welcomes a nearly-increasing amount of teams and participants every year.** The participation has increased in almost every edition of the competition, as shown graphically with the number of participants (red squares) and number of teams (blue circles) registered every year.

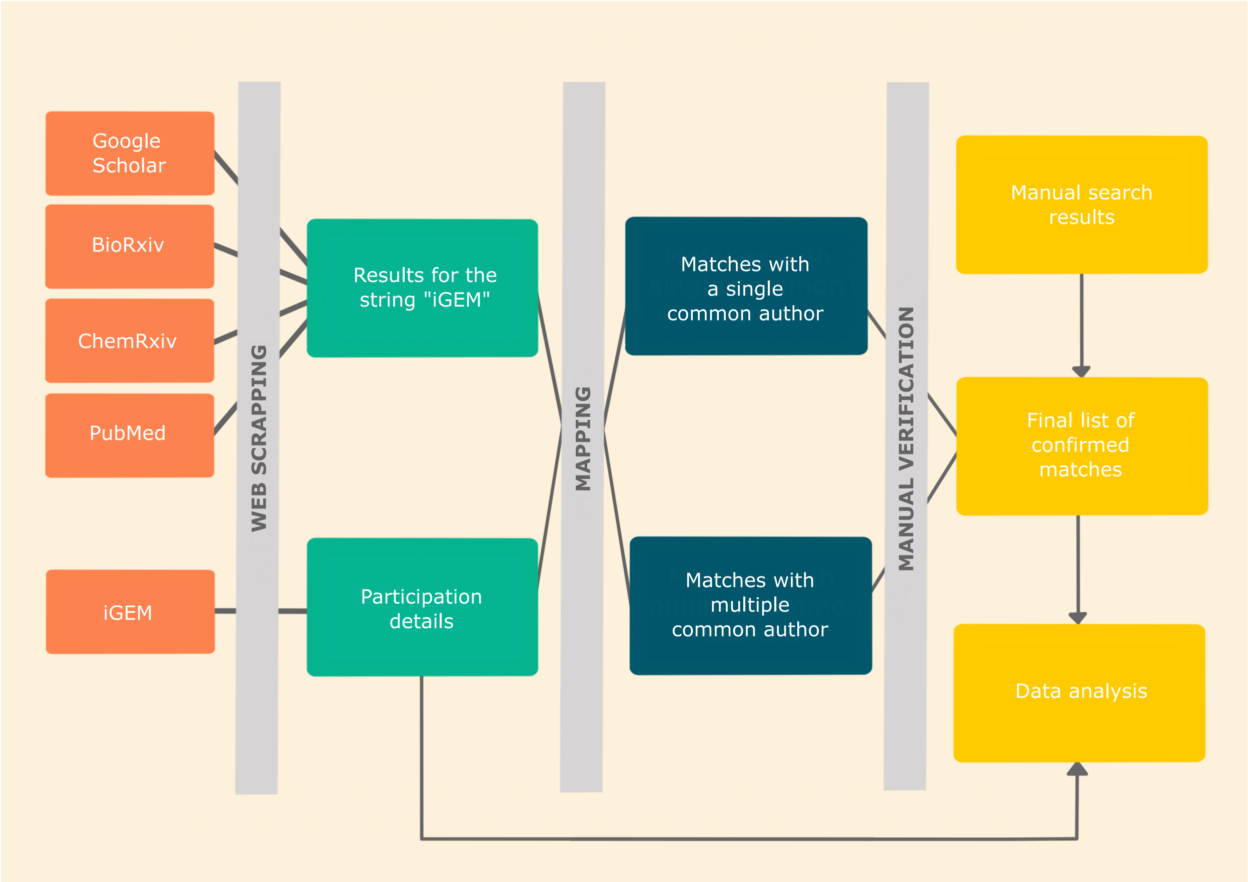


**Figure S2: Web scrapping and mapping pipeline to identify iGEM-derived publications.** Schematic describing the methodology of the pipeline for the identification and matching of iGEM-derived publications. Servers (e.g. Google Scholar, bioRxiv, etc.) are scrapped and correlated with data obtained from www.igem.org by identifying the string "iGEM" in manuscripts. Susequently, the author information is compared with that of the team roster, and common matching authors are used to filter the data. The data set is then verified manually using parameters such as project descriptions and affiliations. The final confirmed list of publications is then analysed for extracting quantitative metrics.

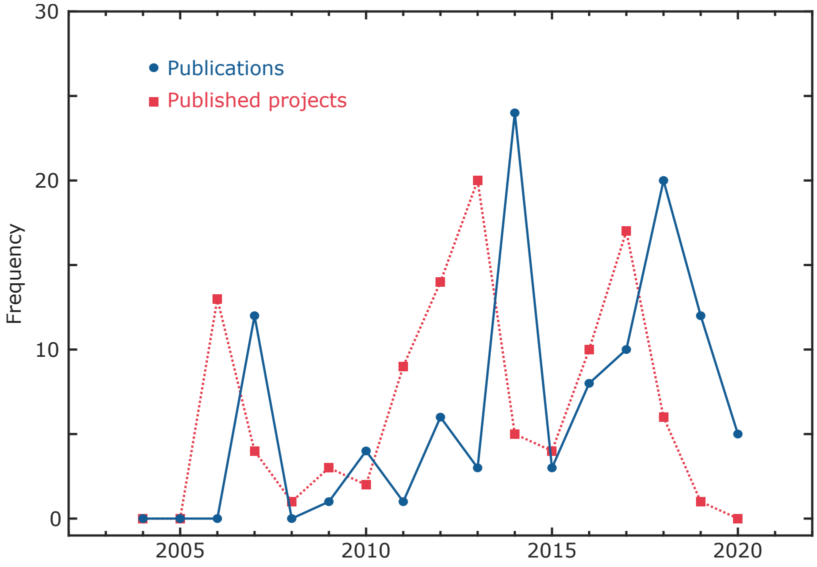


**Figure S3: Frequency of dissemination of iGEM projects in the form of publications.** Trends in the dissemination of iGEM projects published either as preprints or in peer-reviewed venues, conveyed by the number of published teams (red squares) and the number of iGEM-associated publications (blue circles) every year.

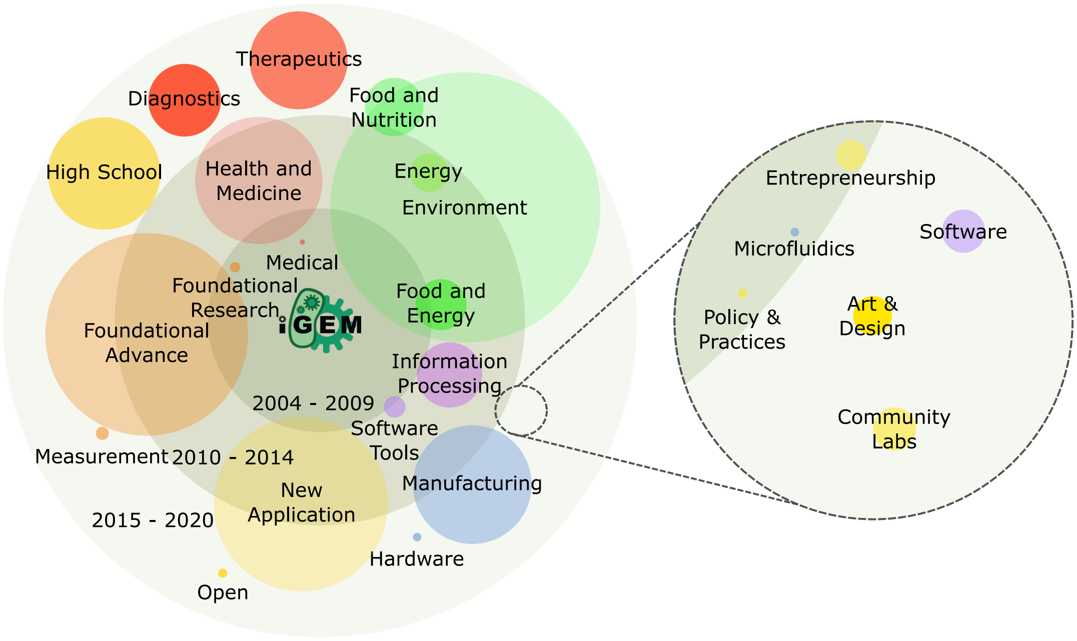


**Figure S4: iGEM projects address a wide range of topics and disciplines.** Bubble chart showing the distribution of the different topics addressed by iGEM projects, as determined by the track each team participated in. In this case, the teams that did not have an assigned track (205) were not included. The radii of the bubbles serve as a proxy for the number of teams in each category, while colour schemes highlight the evolution of the broad topics encompassed by the competition. Concentric circles represent time periods, as denoted by the labels in each of them, which aid in roughly visualising their time association to the competition.

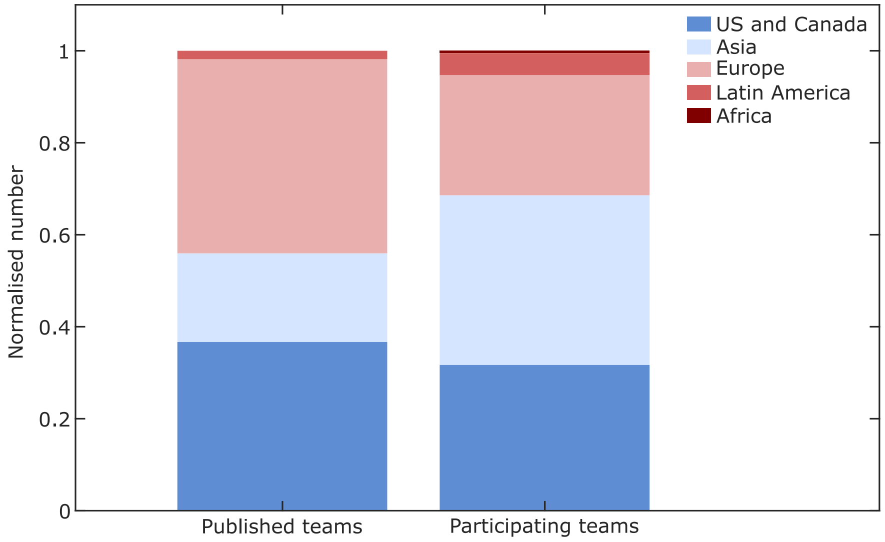


**Figure S5: Normalised regional distribution of publication and participation incidence.** Stacked bar plot showing the publication incidence per region, as defined by iGEM, normalised to the total number of publications (109). For comparison, an analogous bar plot shows the participation incidence per region, normalised to the total number of participating teams (2986).

### Supplementary Tables

**Table S1: iGEM-derived publications disseminated as preprints in the first instance.** Compilation of teams which disseminated their research initially as preprints and subsequently in a peer-reviewed journal. In this case, the Elapsed time ($\Delta t$) is defined as the absolute difference (in months) between the date of publication in the journal and that of the publication in bioRxiv.

| **Team** | **Preprint publication date** | **Journal publication date** | **Elapsed time, $\Delta t$ (months)** |
|---|---|---|---|
| Munich 2017 | July 12, 2019 | December 18, 2019 | 5 |
| William & Mary 2016 | May 09, 2018 | October 29, 2018 | 6 |
| TU-Eindhoven 2018 | August 14, 2019 | February 27, 2020 | 7 |
| Penn 2017 | September 12, 2018 | December 04, 2018 | 3 |
| Arizona State University 2016 | March 09, 2018 | August 23, 2018 | 6 |
